# DPPH-scavenging and antimicrobial activities of Asteraceae medicinal plants on uropathogenic bacteria

**DOI:** 10.1101/2019.12.25.888404

**Authors:** Trinh Canh Phan, Thao Thi Thanh Le, Ha Tran Viet Hoang, TuAnh Nguyen

## Abstract

Asteraceae species were widely applied in traditional medicines in Asia countries as sources of natural antioxidants and antimicrobial agents. This study aimed to evaluate DPPH-scavenging capacities and antimicrobial activities of nine Asteraceae species collected from Southern Vietnam. Antioxidant and antimicrobial activities were determined by standard protocols. Essential oils from *Ageratum conyzoides, Helianthus annuus, Artemisia vulgaris* indicated significant inhibitory effects on *Staphyloccocus aureus* and *Candida* spp.. Crude extracts and fractions from *Taraxacum officinale, Chrysanthemum morifolium, Ageratum conyzoides, Tagetes erecta* showed inhibitory ability on at least one testing bacterial strains including *S. aureus, Escherichia coli, Klebsiella pneumoniae, Pseudomonas aeruginosa*. Study on clinical isolates, ethyl acetate fraction from *A. conyzoides* displayed the most potent effect on uropathogenic *E. coli* and *K. pneumoniae* with MIC at 1.25-10 mg/ml and 5-12.5 mg/ml, respectively. DPPH scavenging assay indicated that *Tagetes erecta* extract had the lowest IC_50_ (17.280 μg/ml) and 2.5 times higher than vitamin C (7.321 μg/ml). This study revealed that *A. conyzoides* has good potential against uropathogenic *E. coli* and *K. pneumoniae*, and could, therefore, apply to prophylactic urinary tract infection.

## 1. Introduction

In recent years, antibiotic resistance has become more sophisticated, putting mankind into the post-antibiotic period. Many clinical *Enterobacteriaceae* strains such as *Escherichia coli* and *Klebsiella pneumoniae* have extended-spectrum beta-lactamase (ESBL) and carbapenem-resistant Enterobacteriaceae (CRE) [1-3]. Especially, polymyxin plasmid-mediated resistance gene (mcr-1) was observed in *E. coli* strains, which was isolated from animals and in patients with infection during 2011-2014, in China [4]. Moreover, it is with profound concern that mcr-1 could be transferred to *K. pneumoniae* and *Pseudomonas aeruginosa* via transformation [4]. In 2016, the first report about mcr-1 gene in a patient with urinary tract infections (UITs) in Pennsylvania, the United States was shown by Jennifer Abbasi [5]. According to recent reports, the causative agents of UITs include uropathogenic *E. coli, Klebsiella pneumoniae, Enterococcus* spp., *Staphylococcus saprophyticus*, group B *Streptococcus* (GBS), *Proteus mirabilis, Pseudomonas aeruginosa, Staphylococcus aureus*, and *Candida spp*. [6-10]. In which, *Escherichia coli* is the most common causative agents of both uncomplicated and complicated UTIs [6].

Herbal extracts and their essential oils have used as foods such as floral teas, beverages, functional foods, bakery products, and traditional medicines in many years, with minimal known “side-effects” on human health. Using herbal remedies might help to reduce dependence on antibiotic therapies and minimizing antibiotic resistance [11].

The Asteraceae family (Compositae) is a widespread family of flowering plants, including 32,913 species names, belonging to 1,911 plant genera, distributed in 13 subfamilies [12]. The tropics, subtropics and temperate regions are the natural habitats of Asteraceae species [13]. They usually contain a large amount of essential oil, polyphenols, flavonoid compounds, which are often studied for antimicrobial and antioxidant activities [14-19].

Although there were many reports for antimicrobial and antioxidant effects of Asteraceae species, applications of these extracts in treating infectious diseases need an evaluation of pathogenic bacterial strains isolated from clinical specimens [11]. In the study, we screened antimicrobial and antioxidant activities of ethanol extracts and essential oils from 9 species of Asteraceae on 30 clinical strains causing urinary tract infection, collected from District 2 Hospital, Ho Chi Minh City, Vietnam. The target was seeking the best extract to apply for a healthcare serum to prevent recurrent UITs. The antioxidant activity might be a protective factor for urinary tract epithelium to avoid impacts of oxidative stress.

## 2. Materials and Methods

We conducted investigations on antimicrobial activities of nine Asteraceae species collected from Southern Vietnam, following Figure 1. In particular, after pre-treating and extracting herbal samples to obtain crude extracts and essential oils, we evaluated the antimicrobial effect by applying the diffusion method. The extracts which show activity were fractionated by n-hexane, chloroform, ethyl acetate, respectively. The well-agar-diffusion method was used to determine the antimicrobial capacities of fractions. Fractions which indicated inhibitory zone were evaluated MIC and MBC. DPPH free-radical scavenging assays were carried out on crude extracts [20, 21].

**Figure 1:**
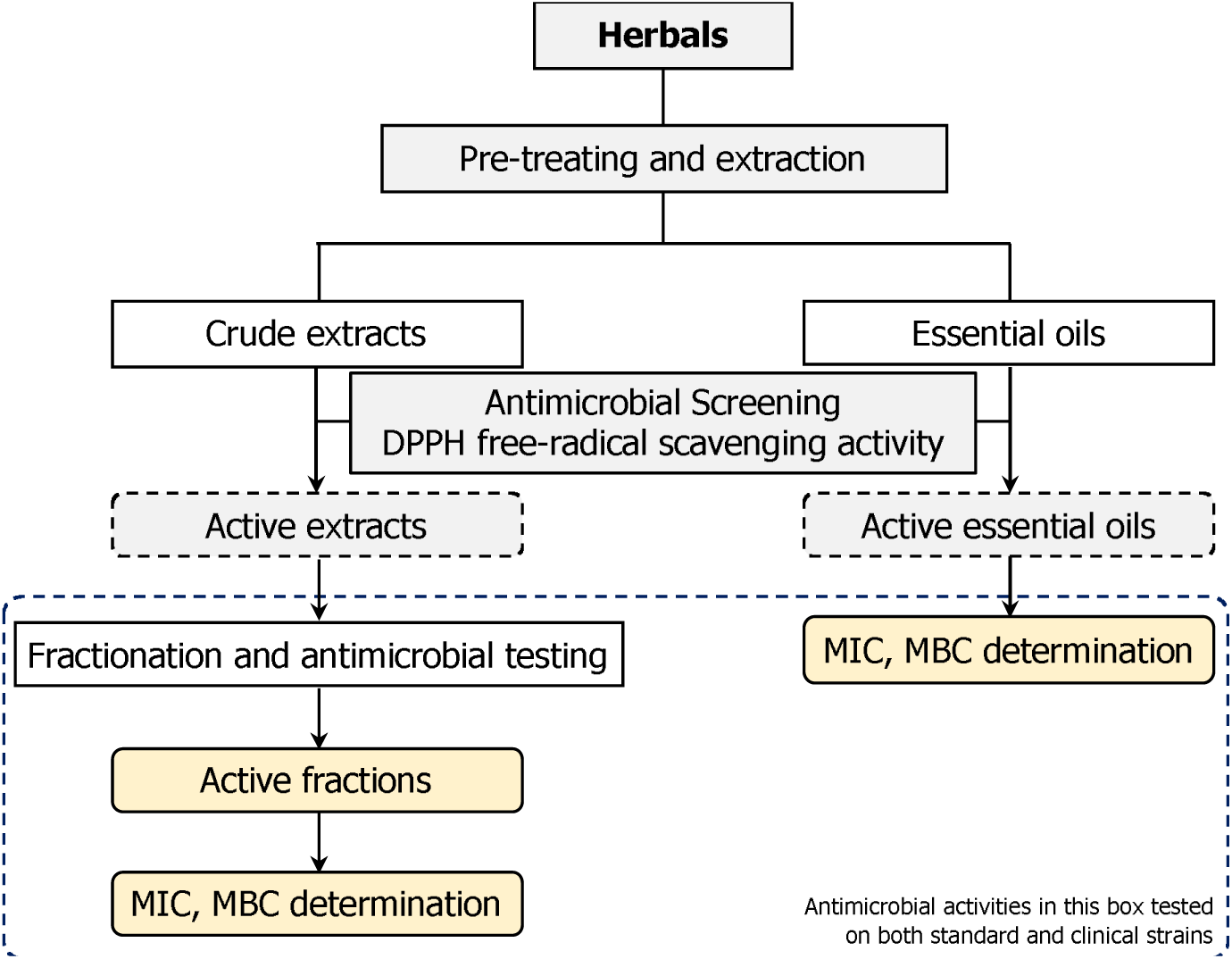
Schematic representation of the experimental layout

### 2.1. Plant Authentication and Preparation

Asteraceae-plant samples were collected from Southern Vietnam from March to May 2016. These samples were identified at the Botany Department, Faculty of Pharmacy, University of Medicine and Pharmacy at Ho Chi Minh City. These species are usually used as traditional medicines in Vietnam. Table 1 shows common names, general uses following the botanical nomenclature of nine species used in this study.

**Table 1:** The investigated plants

| Nomenclature | Common name | Traditional uses* | Part tested<sup>†</sup> |
| --- | --- | --- | --- |
| *Ageratum conyzoides* L. | Billygoat-weed | Sinusitis, anti-inflammation [22]. | Aerial parts, essential oils [22]. |
| *Artemisia vulgaris* L. | Mugwort | Skin ailments, wounds, ulcers [23]. | Aerial parts, essential oils [23]. |
| *Chrysanthemum coronarium* L. | Crown daisy | Pain relief, fever, dysentery [24, 25]. | Aerial parts [25]. |
| *Chrysanthemum morifolium* Ramat. | Florists chrysanthemum | Pimples, dermatitis, fevers [26]. | Flowers [26]. |
| *Helianthus annuus* L. | Sunflower | Anti-inflammatory, malaria [27]. | Flowers, essential oils [27]. |
| *Tagetes erecta* L. | Mexican marigold | Dysentery, asthma, ulcer [28]. | Flowers [28]. |
| *Taraxacum officinale* F. H. Wigg. | Dandelion | Hepatitis, bronchitis, pneumonia [29]. | Aerial parts [29]. |
| *Vernonia amygdalina* Del. | Bitter leaf | Fever, measles, parasites [30]. | Leaves [30]. |
| *Wedelia trilobata* L. | Wedelia | Fever, infection [31]. | Aerial parts [31]. |

\* In this column there are presented only such traditional uses which can imply the presence of antimicrobial compounds. <sup>†</sup> Part tested: the part of the plant used in this study.

After harvesting, the samples were washed under running water to remove dust and rinse with distilled water to drain. Then they were dried in the shade. Afterwards, the dried plant materials were finely grounded by mechanical grinders. The powder was stored in tightly closed glass containers in the dark at room temperature.

### 2.2. Preparation of plant extracts

#### 2.2.1. Preparation of essential oils

Plant samples were cut into small pieces after washing under running water. Plant materials (100 g) were placed in a flask (1 L) together with distilled water. Clevenger apparatus was used for distillation of essential oils. After steam distillation (about 3 hours), the oil was isolated and dried over anhydrous sodium sulfate (Merck). The essential oils were used directly for antimicrobial assay.

#### 2.2.2. Ethanol crude extracts

Plant materials (50 g) were extracted by cold soaking with 500 ml of 96% ethanol (Xilong Chemical) for 24 hours at 10:1 solvent-to-sample ratio (v/w). Then, the mixtures were filtered through Whatman filter paper. The extracts were allowed to evaporate at a temperature of 45-50 °C with water bath WNB 29 (Memmert). These steps were repeated three times to achieve maximal extraction of compounds. Dried crude extracts were weighed and kept at −35 °C till further use (not more than one month). These extracts had been screened antimicrobial activity with well agar diffusion method to choose the good antimicrobial extracts for the next step.

#### 2.2.3. Fractionation of the ethanol crude extracts

Ethanol crude extracts, which possess the strong antimicrobial effect, were carried out liquid/liquid extraction with *n-*hexane, chloroform, and ethyl acetate, respectively. After evaporating solvent, each fraction was determined the antimicrobial activity.

Stock solutions of crude extracts and fractions were prepared at a concentration of 100 mg/ml in 10% dimethyl sulfoxide (DMSO - Merck).

### 2.3. Preliminary phytochemical screening

The ethanol crude extracts were analyzed for phytochemical constituents for the identification of various classes of compounds, according to Maria, Shirley, Xavier, Jaime, David, Rosa and Jodie [32].

### 2.4. Microorganism strains and culture conditions

Microbial strains from American Type Culture Collection (ATCC) were used in this study for preliminary antimicrobial assays, which included methicillin-sensitive *Staphylococcus aureus* ATCC 25923 (SA), methicillin-resistant *S. aureus* ATCC 33591 (SR), *Enterococcus faecalis* ATCC 29212 (EF), *Escherichia coli* ATCC 25922 (EC), *Klebsiella pneumoniae* ATCC 35657 (KP), *Pseudomonas aeruginosa* ATCC 27853 (PA), *Candida albicans* ATCC 10231 (CA). Two clinical *non-albicans* strains being *Candida glabrata* ND31 (CG), *Candida tropicalis* PNT20 (CT), which were provided by Nguyen Dinh Nga et al. (2012) [33]. Clinical isolates (15 *E. coli* and 15 *K. pneumoniae)*, which were isolated from District 2 Hospital, Ho Chi Minh City, Vietnam, in 2016, were also applied for antimicrobial investigation on the potential extracts.

These strains were preserved in 25% glycerol at −80 °C. One strain tube thawed rapidly at 37 °C and cultured in 10 ml Brain heart infusion (BHI - Merck) at 37 °C over 24 hours. The bacteria were streaked on BHI agar (BHA - Merck) at 37 °C over 24 hours. One to five colonies were used to prepare bacterial suspension to match a 0.5 McFarland standard (1-1.5 × 10^8^ CFU/ml). Mueller-Hinton agar medium (MHA - Merck) and MHA supplied 2% glucose medium (MHGA - Merck) were used for determination antibacterial and antifungal activity, respectively.

### 2.5. Antimicrobial diffusion method

The antimicrobial activity of the samples was initially evaluated by well agar diffusion assay for the extracts and disc diffusion assay for the essential oils [21]. The growth medium was poured into Petri dishes at 45-50 °C, approximately 4 mm depth, and they were left to solidify in the laminar-flow hood. Subsequently, a sterile cotton swab was dipped into overnight microbial suspensions (adjusted to a turbidity of 0.5 McFarland standard). Agar plates were inoculated by evenly streaking cotton swab over the agar medium.

As for extracts, wells with a diameter of 6 mm were cut in the inoculated-agar medium with a sterile cork borer. Stock solutions of the samples were diluted in sterile distilled water to get 100 mg/ml concentration. The tested samples and controls (50 μl) were dispensed into the wells.

As for essential oils, 20 μl of the oils were applied on filter paper discs (6 mm). These discs were put on the inoculated-agar surface.

The plates were incubated at 37 °C for 24-48 hours. After that, the diameters of growth inhibition zones were measured. Following control agents were used: positive control agents - ampicillin 20 μg/ ml (for bacteria) and ketoconazole 20 μg/ml (for yeasts); negative control agent - 10% DMSO.

### 2.6. Determination of Minimum Inhibitory Concentration

Determination of minimum inhibitory concentrations (MIC) of the extracts and essential oils was done using agar dilution method [21]. Stock solutions were diluted with melted agar to a concentration range so that the following concentration equal to half the previous concentration. Pour the agar to Petri discs and wait for them to solidify in the laminar-flow hood. Microorganism suspensions at 0.5 McFarland were diluted by 0.85% NaCl solution to reach 1-1.5 × 10^7^ CFU/ml for bacteria and 1-5 × 10^6^ CFU/ml for *Candida* spp.. These suspensions were spotted (1 µl) on the agar surface. Bacterial or yeast colonies growth at the spot after incubating at 37 °C for 24-48 hours indicated for the assay. The MIC is the lowest concentration of antimicrobial agent that completely inhibits growth. The experiment was replicated three times.

### 2.7. Determination of Minimum Bactericidal Concentration

To determine the minimal bactericidal concentration (MBC), the spots at MIC, 2MIC, 4MIC, and 8MIC were washed with 1 ml of 0.85% NaCl. The 100 µl of the washing suspension was spread evenly over BHA agar. After 24-48 hours of incubation at 37 °C, the number of surviving bacteria was determined. The MBC was defined as the lowest extract concentration at which 99.9% of the bacteria have been killed. The experiment was replicated three times.

### 2.8. DPPH free-radical scavenging activity assay

The 0.25 mg/ml 2,2-diphenyl-1-picryl-hydrazyl-hydrate (DPPH – Sigma Aldrich) solution in methanol (working solution) was used to determine the antioxidant capacity. Stock solutions were prepared at the sample concentration of 10 mg/ml. The preliminary tests were carried out on TLC Silica gel 60 F[[[(Merck). After impregnating 2 μl/spot of stock solutions on the TLC plate, the plate was dipped into the DPPH working solution and incubated at the room temperature 30 minutes. The positive test indicated the yellow on the violet background [34].

Reaction mixtures consisted of stock solution, 2 ml DPPH working solution and methanol as a solvent to have a sample concentration from 0.1 to 0.5 mg/ml. The negative controls had only the solvent instead of testing solution. The mixtures were incubated for 30 minutes at 37 °C in the dark. The decrease in the absorbance at 517 nm was measured (A_E_) [34]. The experiment was carried out in triplicate. Samples and positive control ascorbic acid were tested in triplicate over the same range of sample concentrations. Radical scavenging activity was calculated using the following formula:

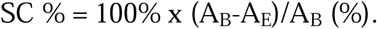

where A_B_: absorbance of the blank sample, and A_E_: absorbance of the plant extract.

The antioxidant activity was expressed as the IC_50_ value. This value was determined from the plotted graphs of scavenging activity against the concentration of the sample.

## 3. Results and Discussion

### 3.1. Results

#### 3.1.1. Preliminary phytochemical screening

Table 2 depicts various classes of phytoconstituents presenting in testing herbs. In general, tannins, flavonoids, phenolics were found in all testing herbs. *A. conyzoides, H. annuus*, and *A. vulgaris* possess a large amount of essential oil.

**Table 2:** Phytochemical profile of ethanol crude extracts

| Group of compounds | TO | CM | AC | CC | HA | VA | AV | TE | WT |
| --- | --- | --- | --- | --- | --- | --- | --- | --- | --- |
| Carotenoid | - | - | - | - | - | - | - | - | - |
| Free triterpenoids | + | - | - | - | - | - | - | + | - |
| Alkaloids | - | - | - | - | + | + | + | + | - |
| Coumarins | - | - | - | - | - | - | - | - | - |
| Anthraglycosides | + | - | - | - | - | + | - | - | - |
| Flavonoids | ++ | +++ | ++ | + | ++ | + | + | +++ | + |
| Heart glycolysis | - | - | - | - | - | - | - | - | - |
| Tannins | +++ | ++ | +++ | + | ++ | ++ | ++ | +++ | ++ |
| Phenolics | +++ | +++ | +++ | ++ | ++ | ++ | ++ | +++ | + |
| Saponins | + | + | - | + | - | + | + | + | + |
| Organic acids | - | - | - | - | - | - | - | - | - |
| Reducing agent | + | + | + | + | + | + | + | + | + |

TO: *T. officinale*; CM: *C. morifolium*; AC: *A. conyzoides*; CC: *C. coronarium*; HA: *H. annuus*; VA: *V. amygdalina*; AV: *A. vulgaris*; TE: *T. erecta*; WT: *W. trilobata*.

#### 3.1.2. Antimicrobial screening assay

Among ethanol crude extracts tested, there were four plant species including *Ageratum conyzoides, Chrysanthemum morifolium, Tagetes erecta*, and *Taraxacum officinale* indicated the antibacterial activities from one to four standard bacterial strains. None of them had the anti-yeast activity on three yeast strains listed in Table 3. To be more specific, *A. conyzoides* and *T. erecta* expressed the broad-spectrum antimicrobial effect on Gram positive (20-22 mm on MSSA and MRSA) and Gram negative bacteria (11-18 mm), meanwhile *T. erecta* the inhibitory capacity on *P. aeruginosa* (13 mm). *C. morifolium* and *T. officinale* indicated the activity only on methicillin-sensitive *S. aureus*.

**Table 3:** The inhibitory zone diameter (IZD, mm) of the ethanol crude plant extracts determined by agar well diffusion assay

| Plant species | Inhibitory Zone Diameter (mm) |  |  |  |  |  |  |  |  |
| --- | --- | --- | --- | --- | --- | --- | --- | --- | --- |
|  | SA | SR | EF | KP | EC | PA | CA | CG | CT |
| *A. conyzoides* | 22 | 20 | - | 18 | 11 | - | - | - | - |
| *T. erecta* | 21 | 20 | - | 16 | - | 13 | - | - | - |
| *C. morifolium* | 14 | - | - | - | - | - | - | - | - |
| *T. officinale* | 10 | - | - | - | - | - | - | - | - |
| *A. vulgaris* | - | - | - | - | - | - | - | - | - |
| *C. coronarium* | - | - | - | - | - | - | - | - | - |
| *H. annuus* | - | - | - | - | - | - | - | - | - |
| *V. amygdalina* | - | - | - | - | - | - | - | - | - |
| *W. trilobata* | - | - | - | - | - | - | - | - | - |

- no inhibitory zone

The ethanol crude extracts, which showed antimicrobial activity (active extracts), were decanted with different polarization solvents included *n*-hexane, chloroform, ethyl acetate to obtain fractions. Diffusion method was conducted to determine antimicrobial activities of these fractions on microbial strains. Noticeably, antimicrobial agents usually distribute in moderate polarity fractions such as chloroform and ethyl acetate (Table 4). The ethyl acetate fraction of *A. conyzoides* displayed the IZD on MSSA (23 mm), MRSA (21 mm), *K. pneumoniae* (15 mm), and *E. coli* (14 mm). Similarly, *T. erecta’s* fraction of ethyl acetate shows IZD from 11-18 mm on MSSA, MRSA, *K. pneumoniae* and *P. aeruginosa*.

**Table 4:** IZD (mm) of the fractions determined by well agar diffusion assay

| Plant species | Part tested | Fractions | IZD (mm) |  |  |  |  |
| --- | --- | --- | --- | --- | --- | --- | --- |
|  |  |  | SA | SR | KP | EC | PA |
| *A. conyzoides* | Aerial parts | *n* – hexane | - | - | - | - | - |
|  |  | CHCl<sub>3</sub> | 11 | 11 | - | - | - |
|  |  | EtOAc | 23 | 21 | 15 | 14 | - |
|  |  | EtOH | 9 | - | 10 | - | - |
| *T. erecta* | Flower | *n* – hexane | 11 | - | 11 | - | - |
|  |  | CHCl<sub>3</sub> | 11 | - | 10 | - | - |
|  |  | EtOAc | 17 | 12 | 18 | - | 11 |
|  |  | EtOH | - | - | - | - | - |
| *C. morifolium* | Flower | *n* – hexane | - | - | - | - | - |
|  |  | CHCl<sub>3</sub> | 13 | - | - | - | - |
|  |  | EtOAc | 14 | - | - | - | - |
|  |  | EtOH | - | - | - | - | - |
| *T. officinale* | Aerial parts | *n* – hexane | - | - | - | - | - |
|  |  | CHCl<sub>3</sub> | - | - | - | - | - |
|  |  | EtOAc | 11 | - | - | - | - |
|  |  | EtOH | - | - | - | - | - |

- no inhibitory zone

The essential oils of *A. conyzoides* (aerial parts), *A. vulgaris* (aerial parts), and *H. annuus* (flowers) were demonstrated the antimicrobial activity (Table 5). Although ethanol crude extracts of *A. vulgaris* and *H. annuus* do not have inhibitory capacity on testing microorganisms, this test reveals the effect of their essential oils on *S. aureus* and *Candida* spp.

**Table 5:** Antimicrobial activity of the essential oils determined by disc diffusion assay

| Essential oils | IZD (mm) |  |  |  |  |  |  |  |  |
| --- | --- | --- | --- | --- | --- | --- | --- | --- | --- |
|  | SA | SR | EF | KP | EC | PA | CA | CG | CT |
| *A. conyzoides* | 15 | - | - | - | - | - | - | 8 | 10 |
| *A. vulgaris* | 23 | 10 | - | - | - | - | 14 | 15 | 16 |
| *H. annuus* | 10 | - | - | - | - | - | - | - | 11 |

- no inhibitory zone

#### 3.1.3. MIC values of active extracts and essential oils on standard strains

MIC and MBC values of the selected four plant parts: *Ageratum conyzoides, Chrysanthemum morifolium, Tagetes erecta*, and *Taraxacum officinale* crude extracted, and four fractions were determined (Table 6).

**Table 6:**
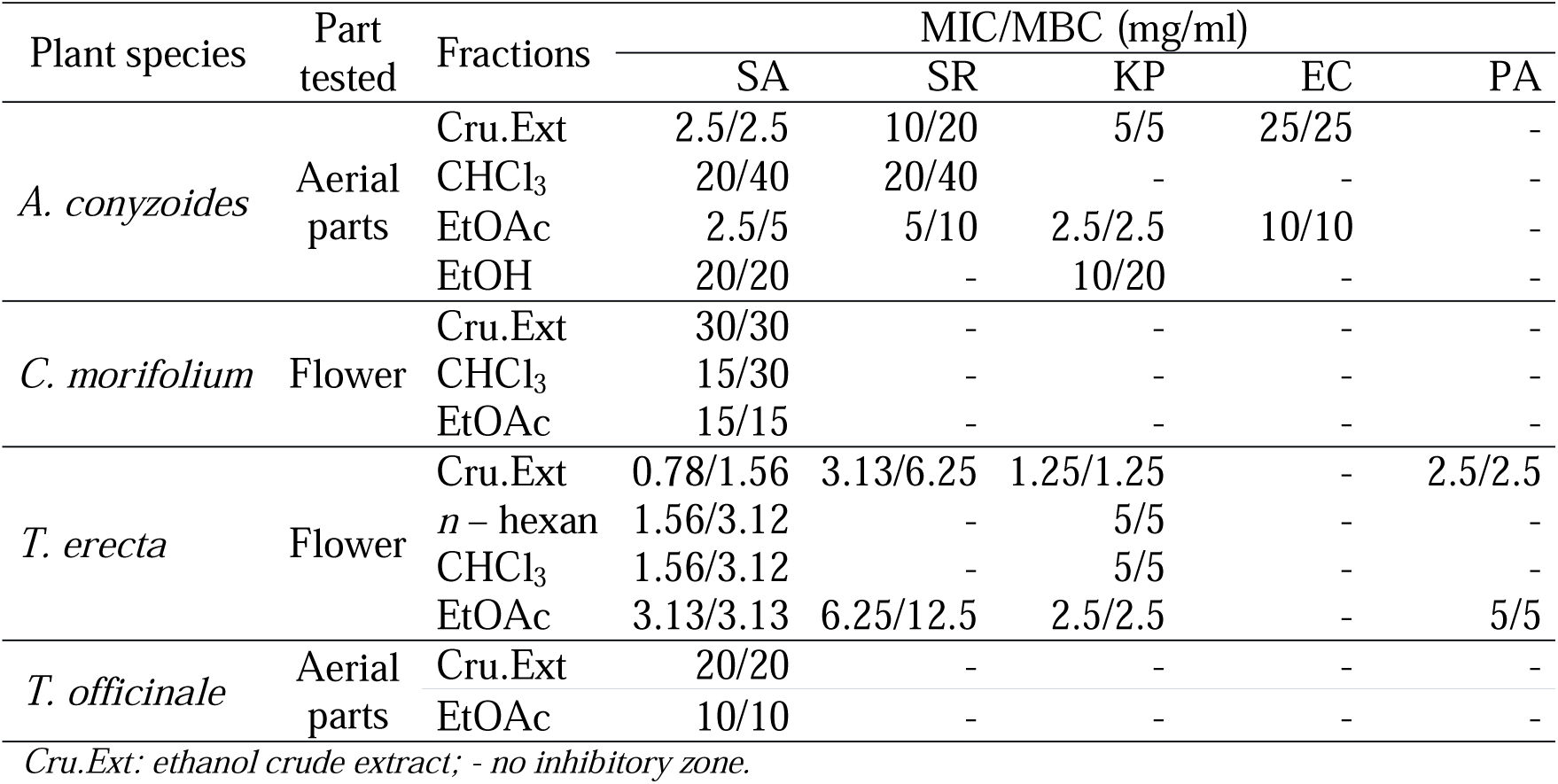
MIC and MBC values (mg/ml) of crude extracts and fractions of four selected plant materials

MIC and MBC values of the three essential oils, *A. conyzoides, A. vulgaris*, and *H. annuus*, were shown in Table 7.

**Table 7:** Determination of minimum inhibitory concentration (MIC) of essential oils

| Essential oils | MIC/MBC (μl/ml) |  |  |  |  |
| --- | --- | --- | --- | --- | --- |
|  | SA | SR | CA | CG | CT |
| *A. conyzoides* | 3.75/7.50 | - | - | 7.5/15 | 10/20 |
| *A. vulgaris* | 2.50/2.50 | 6.25/6.25 | 6.25/12.5 | 5/10 | 5/10 |
| *H. annuus* | 3.75/7.50 | - | - | - | 10/20 |

#### 3.1.4. Antimicrobial activity on uropathogenic strains of *E. coli* and *K. pneumoniae*

Through these results, we found the anti-Gram-negative-bacteria effect of extracts from *A. conyzoides* and *T. erecta* on *K. pneumoniae* and *E. coli* being two main pathogenic bacteria of urinary tract infection. In order to explore the best extract to prevent recurrence UITs, we evaluated the crude extracts and fractions of *A. conyzoides* and *T. erecta* on 15 *E. coli* and 15 *K. pneumoniae* isolates from urine specimens at District 2 Hospital, Ho Chi Minh city.

The well agar diffusion assay was used for analyzing the activity of the investigated extracts. The results showed that only the ethyl acetate fraction from *A. conyzoides* had antibacterial activity against tested isolates. MIC values of the ethyl acetate fraction of *A. conyzoides* were determined by agar dilution assay on the isolates of *E. coli* and *K. pneumoniae* that had minimum and maximum diameters of growth inhibition zone as well as on ESBL-producing isolates. The width of inhibition zones and minimum inhibitory concentration (MIC) showed in Table 8. Specifically, the IZD on *E. coli* and *K. pneumoniae* indicated from 10.83 mm (E3) to 23.72 mm (E72), and from 9.33 mm (K18) to 19.73 mm (K17), respectively. *A. conyzoides’s* ethyl acetate fraction displayed MIC values on E3, E72, K17, and K18 being 12.5, 5, 10, and 1.25 mg/ml, respectively. ESBL-producing strains, E68 and K26 expressed MIC 6.25 and 2.5 mg/ml, respectively.

**Table 8.** Antimicrobial activity of the ethyl acetate fraction of *A. conyzoides* determined by agar well diffusion assay

| Strains | IZD (mm) | Strains | IZD (mm) |
| --- | --- | --- | --- |
| E3 | 10.83 | K18 | 9.33 |
| E7 | 11.43 | K27 | 10.03 |
| E25 | 12.17 | K19 | 11.23 |
| E36 | 13.33 | K20 | 11.37 |
| E77 | 13.43 | K15 | 11.67 |
| E38 | 13.67 | K23 | 11.83 |
| E27 | 14.47 | K14 | 11.93 |
| E39 | 14.47 | K21 | 12.10 |
| E84 | 14.97 | K28 | 12.37 |
| E68* | 15.43 | K25 | 13.23 |
| E63 | 20.37 | K22 | 13.27 |
| E51 | 21.40 | K16 | 14.20 |
| E42 | 22.37 | K29 | 17.67 |
| E94 | 22.37 | K26* | 19.27 |
| E72 | 23.27 | K17 | 19.73 |

\* ESBL-producing strain

#### 3.1.5. Free radical scavenging activity

DPPH screening test showed the good antioxidant effect of all crude extracts and essential oils. Free radical scavenging activity of total ethanol extracts was quantitatively determined using a DPPH assay. IC_50_ value represents the concentration of tested extract, where the inhibition of test activity reached 50% (Figure 2). The results were graphed by Microsoft Excel 2016. The IC_50_ values of essential oils were too large (3000 - 6000 μg/ml), which were outside the linear range.

**Figure 2:**
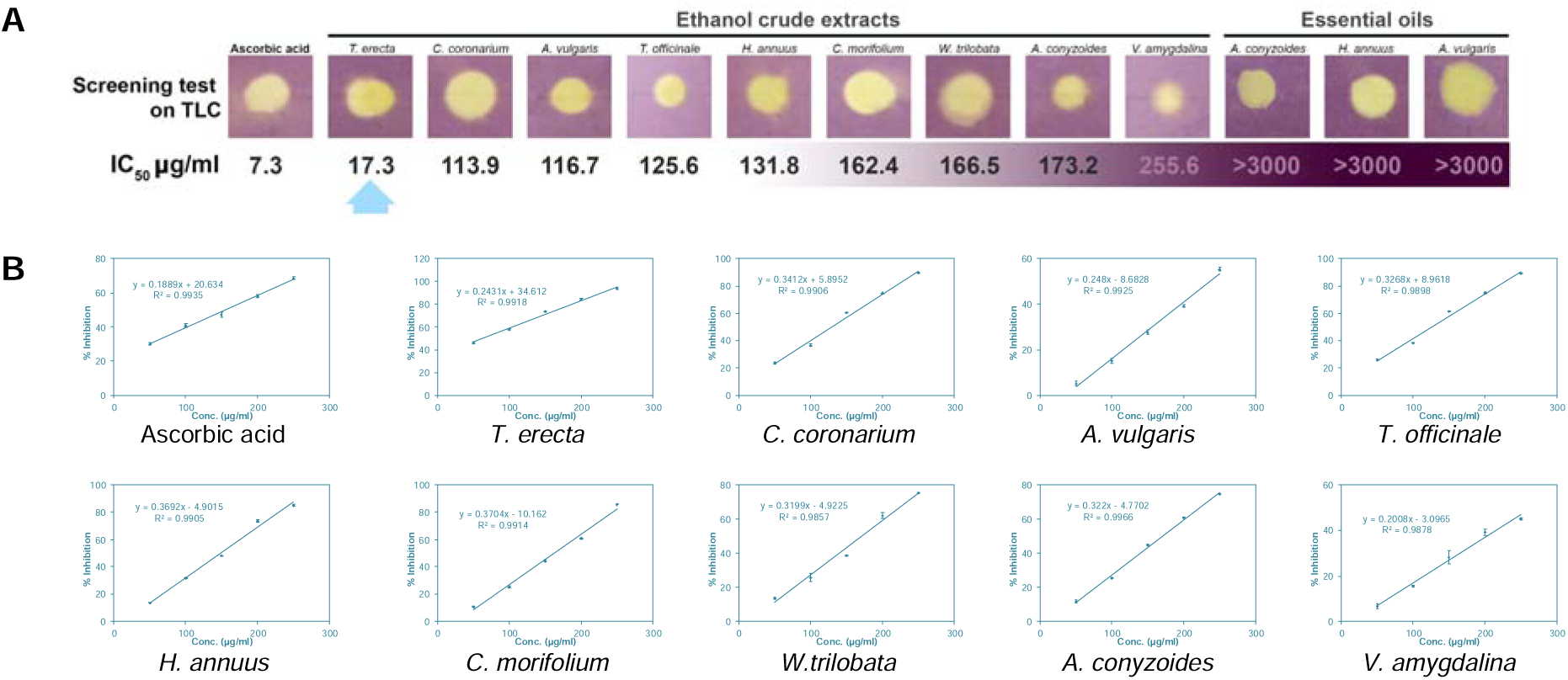
(A) DPPH screening test on TLC and IC_50_ values. (B) The plotted graphs of scavenging activity against the concentration of the ethanol crude extracts and ascorbic acid

*T. erecta* shows the significant scavenging effect for free radicals with IC_50_ = 17.3 µg/ml, 2.4 folds comparing to ascorbic acid.

### 3.2. Discussions

Among the tested extracts and essential oils, there were four crude extracts (*Ageratum conyzoides, Chrysanthemum morifolium, Tagetes erecta*, and *Taraxacum officinale)* and three essential oils (*A. conyzoides, A. vulgaris*, and *H. annuus*) indicated the antimicrobial activity against nine bacteria and three yeast strains. While the crude extracts only had effects on bacteria, the essential oils had effects on both bacteria and yeast strains. There are many reports about antiseptic, antimicrobial, antioxidant, and insecticidal activities of essential oils [35]. Chemical constituents represent in essential oils usually derived from terpenes, phenolic compounds, and aromatic or aliphatic acid esters, which can partition into the lipids of bacterial and mitochondrial membrane resulting in disturbing the cell structures. The death of cell caused by leakage of a large number of essential molecules and ions from the bacterial cell [36].

In this research, the ethanol-crude-extract from *Tagetes erecta* flower showed the highest inhibitory effect on MSSA (0,78 mg/ml), MRSA (3.13 mg/ml), *K. pneumoniae* (1.25 mg/ml), and *P. aeruginosa* (2.5 mg/ml), in comparison with other extracts. Recently, many reports demonstrated that the extracts of *Tagetes erecta* inhibited *S. aureus, P. aeruginosa, K. pneumoniae*, and *P. mirabilis* [37-39]. Padalia and Chanda [37] reported MIC values on those bacteria in the range of 62-1250 µg/ml.

*Ageratum conyzoides* aerial part extracts had an effect on MSSA, MRSA, *K. pneumoniae*, and *E. coli*, but the MIC values were higher than *T. erecta* extracts. Amadi, Oyeka, Onyeagba, Ugbogu and Okoli [40] demonstrated ethanol extract of *A. conyzoides* shows the significant inhibitory zone on *Streptococcus mutans*. Following the report of Akinyemi, Oladapo, Okwara, Ibe and Fasure [41], ethanol extract from *A. conyzoides* indicated MIC and MBC on MRSA of 43 µg/ml and 63.2 µg/ml, respectively. Kouame, Toure, Kablan, Bedi, Tea, Robins, Chalchat and Tonzibo [42] studied on essential oils extracted from flower and stem of *A. conyzoides*; the results displayed antibacterial activities with MIC in the range of 64 to 256 µg/ml on both Gram-positive and Gram-negative bacteria [42]. In this study, the MIC values were higher than previous reports [41, 42]; these differences can relate to the dissimilarity of the time of sampling and extraction condition.

Ethyl acetate fractions (EA) from *A. conyzoides, T. erecta, C. morifolium*, and *T. officinale* showed activities against *S. aureus*. Noticeably, EA from *A. conyzoides* and *T. erecta* displayed considerable impacts on Gram-negative bacteria including *E. coli, K. pneumoniae* (for *A. conyzoides*), and *P. aeruginosa* (for *T. erecta*). However, evaluating these EA on uropathogenic isolates, there was only the EA of *Ageratum conyzoides* expressed the capacity against uropathogenic isolates. Notably, it was active against *E. coli* and *K. pneumoniae* producing ESBL – the strains that were highly resistant in clinical infection. Hence, *Ageratum conyzoides* is a good candidate for anti-uropathogenic bacteria.

*A. conyzoides, A. vulgaris*, and *H. annuus* essential oils expressed antimicrobial activities against both bacteria (MSSA, MRSA) and yeasts (*C. albicans, C. glabrata, C. tropicalis*). Our investigations witnessed anti-MRSA and anti-*Candida* effects of *A. vulgaris* essential oil. Those capacities could be attributed to a large amount of mono- and sesquiterpenes compounds such as sabinene, β-thujone, chrysanthenone, camphor, borneol in *A. vulgaris* oil [43].

DPPH is a free radical, stable at room temperature, which produces a violet solution in methanol. It is reduced in the presence of an antioxidant molecule, making the color of the solution turned yellow. The use of DPPH provides an easy and rapid way to evaluate antioxidants.

All nine ethanol extracts were capable of capturing DPPH free radicals. *T. erecta* extracts had the lowest IC_50_ value, 17.3 µg/ml, which is only 2.5 times higher than the IC_50_ value of ascorbic acid. This result could be due to the presence of a large amount of flavonoid and phenolic compounds in *T. erecta* as mentioned in preliminary phytochemicals. In previous studies, marigolds have shown to contain quercetagetin, glucoside of quercetagetin, phenolics, syringic acid, methyl 3,5- dihydroxy-4-methoxy benzoate, quercetin, thienyl, and ethyl gallate [44]. Quercetin and ethyl gallate are potent antioxidant compounds in both *in vivo* and *in vitro* [45-47]. Containing many compounds with strong free radical scavenging effects of *T. erecta* showed the ability for treatment of diseases caused by free radicals such as cancer, diabetes, cardiovascular, etc. However, this capacity needs to be evaluated more accurately by *in vivo* tests.

## 4. Conclusions

The fraction of ethyl acetate extracted from *A. conyzoides* possesses antimicrobial activities on uropathogenic *E. coli* and *K. pneumoniae* collected from District 2 Hospital, Ho Chi Minh City, Vietnam. This fraction is the potential to apply for healthcare serum in prophylactic recurrence UITs. *T. erecta* showed the highest potent in DPPH radical scavenging assay, which could be become a good candidate for the anti-oxidative agent in food and cosmetic products.

## 5. Data Availability

The data that support the findings of this study are available from the corresponding author upon reasonable request.

## 6. Conflicts of Interest

The authors declare that there are no conflicts of interest regarding the publication of this paper.

## 7. Funding Statement

This study was financially supported by Univerisity of Medicine and Pharmacy at Ho Chi Minh City, through the grant No. 240-17.

## 8. Acknowledgements

The authors would like to express their gratitude to Associate Professor Nguyen Dinh Nga for providing the clinical microorganisms.

## Notes

### Competing Interest Statement

The authors have declared no competing interest.

### Summary of Updates

revised version after peer review

